# The Driving W Hypothesis as an Explanation for Low Within-Population Mitochondrial DNA Diversity and Between-Population Mitochondrial Transfer

**DOI:** 10.1101/2025.05.31.656024

**Authors:** Darren Irwin

**Affiliations:** Department of Zoology, and Biodiversity Research Centre, University of British Columbia, Vancouver, Canada

**Keywords:** centromere drive, mitonuclear discordance, selective sweep, sex chromosome, W chromosome, Z chromosome

## Abstract

The fields of evolutionary biology, molecular ecology, genetics, and taxonomy have been profoundly influenced by studies of variation in mitochondrial DNA (mtDNA), yet there are often surprising differences between mtDNA and nuclear DNA in the within- and between-population relationships that they display. Here I articulate and evaluate a hypothesis that may explain many of the cases in which mtDNA shows little within-population variation and recent movement between populations. Many taxonomic groups (e.g., birds, butterflies and moths; most snakes; some amphibians, fish, and plants) have sex chromosome systems in which females are heterogametic (i.e., ZW females and ZZ males). If a W chromosome undergoes a mutation that gives it a transmission advantage in getting into the one egg produced by female meiosis, it will tend to cause a female-biased sex ratio in the offspring of females that carry that driving W chromosome. This sex ratio bias increases the frequency of the driving W in relation to the non-driving W in the next generation. In the great majority of species in which mitochondria are inherited matrilineally, the spread of the driving W through the population will carry along the particular mitochondrial genome that happens to be associated with the driving W. I summarize evidence in support of the seven components of this W-mtDNA Drive Hypothesis and present simulations and mathematical formulae showing that W drivers spread much more rapidly than equivalent-strength Z or autosomal drivers. Suppressors of W drive spread at a vastly lower rate. I conclude that many cases of low within-population mitochondrial diversity and mitochondrial transfer between species might be explained by the spread of driving W chromosomes.

## Introduction

Mitochondrial DNA (mtDNA) has played a central role in the development of the fields of molecular population genetics, taxonomy, biogeography, and speciation of eukaryotes (Avise, 2000; Ballard & Rand, 2005; Ballard & Whitlock, 2004; Dowling et al., 2008; Hebert et al., 2003; Hill, 2019; Tobler et al., 2019; Wüster, 2025). This is largely due to its high copy number within cells and its typically uniparental inheritance, making the production of clean sequence more technically feasible compared to nuclear DNA. Uniparental and largely non-recombining inheritance also means that evolutionary relationships of mtDNA sequences are well represented by bifurcating phylogenies (Avise et al., 1987). This combined with an assumption of neutrality has led to the development of coalescence theory to explain and interpret patterns of mtDNA variation within and between species (Kingman, 1982a, 1982b; Wakeley, 2008), with expected mathematical relationships connecting within-group mtDNA variation and effective population size. Often, surveys of mtDNA have been useful in clarifying genetic groups that were predicted based on morphological variation and in estimating divergence times between such groups (Avise, 2000; Hebert et al., 2003). Mitochondrial DNA data provided some of the earliest data on genetic variation within species, sometimes revealing striking geographic structure that inspired the invention of the term “phylogeography” (Avise et al., 1987).

Yet mtDNA variation has shown some surprising patterns, leading to vigorous debate regarding the explanations for these patterns as well as the proper role for mitochondrial data in taxonomy, conservation, and evolutionary inference (Ballard & Whitlock, 2004; Bensch et al., 2006; S. Edwards & Bensch, 2009; G. D. D. Hurst & Jiggins, 2005; Irwin, 2002, 2012; Rubinoff & Holland, 2005; Toews & Brelsford, 2012; Zink & Barrowclough, 2008). There are many cases of mitochondrial phylogeographic patterns that were not well predicted based on subspecies descriptions or morphological groups (e.g., Dias et al., 2018; Funk & Omland, 2003; Toews & Brelsford, 2012). Surprisingly little mtDNA variation is sometimes seen across the range of a species in which there are known morphological groups (Ball & Avise, 1992). When phylogeographic structure in mtDNA is observed, there is sometimes extremely low within-group variation compared to the evolutionary distance between the groups (e.g., Toews & Irwin, 2008). Comparisons with nuclear DNA often reveal strong discordance (often referred to as “mitonuclear discordance”): one of these can show much stronger geographic structure than the other, and when both show structure there is often evidence for mtDNA and nuclear DNA showing different levels of gene flow across transition zones between geographic regions (Lee-Yaw et al., 2014; Toews et al., 2014; Toews & Brelsford, 2012). In extreme cases, groups that differ strongly in nuclear DNA can have virtually identical mitochondrial sequences, an observation that has been explained as “mitochondrial capture”—the rapid spread of a mitochondrial clade between two otherwise distinct populations (Ferreira et al., 2018; Irwin et al., 2009; McCallum et al., 2024; Read et al., 2025; Taylor et al., 2021; Weckstein et al., 2001; Wilson & Bernatchez, 1998).

Explanations in the literature for some of these patterns have centred on the concepts of effective population size, genetic drift, sex-biased dispersal, demographic effects, parasite infections that skew the sex ratio, and natural selection on the mitochondria itself (Ballard & Whitlock, 2004; Berlin et al., 2007; G. D. D. Hurst & Jiggins, 2005; Toews & Brelsford, 2012). Because of its matrilineal inheritance (in the great majority of cases), each generation mtDNA has ¼ the possible pathways of inheritance compared to autosomal DNA (assuming distinct sexes rather than hermaphroditic individuals). The consequence is that in an idealized population of female and male individuals (randomly mating, constant population size, no selection), the effects of genetic drift alone cause the expected genealogical coalescence time of mtDNA to be ¼ that of nuclear DNA. The drift process is thereby expected to cause two populations to become reciprocally monophyletic in mtDNA faster than in a segment of nuclear DNA. Add to this the potential for recombination and independent segregation in nuclear DNA, and the net result is less expected phylogeographic structure in nuclear DNA than in mtDNA. Because mtDNA is one large linked sequence, any chance effect (e.g., rare hybridization and backcrossing followed by expansion in the new population due to drift) applies to the entire mitochondrial genome, potentially contributing to cases of mitochondrial introgression across hybrid zones. Sex-biased dispersal has been another invoked explanation for different levels of introgression between mitochondrial and nuclear DNA, as larger dispersal distances of females would tend to spread mtDNA more than autosomal DNA, and the reverse with larger dispersal distances of males (Petit & Excoffier, 2009; Rheindt & Edwards, 2011). Demographic effects can contribute to mito-nuclear discordance as well—for instance population bottlenecks followed by expansions can affect mitochondrial and nuclear DNA diversity differently due to their different means of inheritance. Finally, mtDNA variation could be shaped by selective forces (reviewed by Dowling et al., 2008), either directly via fitness differences due to the mitochondrial sequences (Hill, 2016, 2019; Toews et al., 2014), or indirectly via fitness variation due to other genetic elements that are also matrilineally inherited, such as reproductive parasites (e.g, *Wolbachia*; Gaunet et al., 2019; G. D. D. Hurst & Jiggins, 2005) or female-limited sex chromosomes. These selective forces have the potential to cause low levels of within-population variation, and they each can lead to selective sweeps between populations. In evolutionary groups with ZW sex chromosomes, the mitochondria and the W chromosome are both inherited matrilineally and are thereby genealogically linked (Berlin & Ellegren, 2001; Cosmides & Tooby, 1981). Any process affecting the tendency of a W variant to spread will simultaneously affect the mitochondrion it occurs with (and vice versa).

Emphasizing this matrilineal genealogical linkage, Berlin et al. (2007) hypothesized that there is often diversity-reducing selection on the W chromosome, resulting in reduced within-population diversity in the mtDNA. As W chromosomes usually have a large non-recombining region, linked selection on multiple deleterious and/or advantageous mutations can result in reduced diversity over the whole region (the “Hill-Robertson effect”). Based on this hypothesis, Berlin et al. predicted that species with ZW sex chomosomes (heterogametic females) would tend to have lower mtDNA diversity than species with XY chromosomes (heterogametic males). They found that average mtDNA synonymous site diversity is 0.027 in bird species (ZW) compared to the much larger value of 0.086 in mammal species (XY), about a three-fold difference. They tentatively concluded that “the finding that birds have lower mtDNA variability than mammals is most easily explained by Hill–Robertson effects acting on the W chromosome” and recommended much more analysis involving other taxonomic groups and approaches. Responding to this conclusion, Lane (2008) argued that low variation in mtDNA and W chromosomes is more likely to be due to selection on the mitochondrial genome, given that it encodes proteins that are central to metabolic function—something that might be particularly important in birds due to the metabolic demands of flight. Both Berlin et al. (2007) and Lane (2008) are likely to be at least partially correct, as selection undoubtedly occurs on both the mitochondrial genome and W chromosomes and their genealogical linkage means that selection on either can reduce diversity in both.

In this article, I articulate and explore an additional hypothesis that may go a long way in explaining some of the peculiarities of mtDNA variation and is related to the genealogical linkage to W chromosomes. The term “meiotic drive” refers to an advantage of a certain type of allele in being transmitted from heterozygous individuals into their gametes (Clark & Akera, 2021; F. Finseth, 2023; Henikoff et al., 2001; Kursel & Malik, 2018; Malik & Bayes, 2006; Sandler & Novitski, 1957; Searle & de Villena, 2022; Searle & Pardo-Manuel de Villena, 2024; Talbert & Henikoff, 2022). There are many known empirical examples of meiotic drive, but mostly in the context of autosomes or XY sex chromosome systems (Baird et al., 2023; Jaenike, 2001; Jensen et al., 2024; Lindholm et al., 2016). In male meiosis, four sperm are produced during meiosis of a single pre-meiotic cell; driving alleles cause reduced success of sperm without the driving allele (hence the term “sperm-killing meiotic driver”). In female meiosis, a single egg is produced from a single pre-meiotic cell, such that alleles that are particularly good at getting into the egg have a transmission advantage. In the context of a ZW sex chromosome system, if a variant of the W chromosome has an advantage in being transmitted to eggs then it will tend to spread through a population, and the mtDNA that happens to occur with it will also spread. Repeated cycles of this process will keep within-population mtDNA diversity low and could occasionally lead to introgressive sweeps between populations. This hypothesis is related to but distinct from the Berlin et al. (2007) idea that selective interactions on the W can shape variation in mtDNA, for two reasons: First, Berlin et al. (2007) did not mention meiotic drive. Second, Berlin et al. focused on interactions among selected loci (i.e., Hill-Robertson effects) on the same DNA strand, and meiotic drive can be due to essentially a single locus (i.e., the centromere). Furthermore, selection for a genetic element to get into a successful gamete is often thought of as a distinct process compared to selection based on whole-organism fitness.

We can refer to this hypothesis as the Driving W Hypothesis for Low Mitochondrial DNA Diversity, or more briefly, the W-mtDNA Drive Hypothesis. Some elements of this hypothesis have been examined in previous work: the potential for centromeric regions of chromosomes to experience mutations that give them a transmission advantage has been widely discussed (e.g., Clark & Akera, 2021; Henikoff et al., 2001), and the genealogical connection between female-limited sex chromosomes and matrilineally-inherited cytoplasmic elements has also been emphasized by some researchers (Cosmides & Tooby, 1981). However, I am not aware of a prior clear articulation of the idea that recurrent phases of meiotic drive of the W can cause a widespread pattern of reduced mtDNA variation in species with ZW sex determination. Hence my goal in this paper is to provide a clear summary of the logical components of this hypothesis, summarize evidence for their plausibility, and show simulations and mathematical formulae that demonstrate the particular effectiveness of W drivers compared to Z and autosomal drivers. Furthermore, I evaluate the potential for suppressors of meiotic drive to limit the spread of W drivers.

### Support for seven components of the hypothesis

The logic underlying the Driving W Hypothesis for Low Mitochondrial DNA Diversity can be broken down into seven simple propositions as follows, each motivated by evidence that I summarize further below:

1. **In the great majority of eukaryotes, mtDNA is inherited almost entirely from the female parent, such that there is not recombination between divergent types.**
2. **In taxonomic groups that have ZW sex chromosomes (meaning females are heterogametic ZW and males are homogametic ZZ), the W chromosome is inherited from the female parent.**
3. **In groups where both #1 and #2 apply, mtDNA and the W chromosome are inherited from the same parent, and thereby genealogically linked.**
4. **In both animals and plants, female meiosis produces one viable haploid reproductive cell from two asymmetric cell divisions (meiosis I and meiosis II), and there is a difference in the spindle fibers that do or do not lead toward that viable cell (Fig. 1A).**
5. **Chromosomes can vary in their ability to interact with the spindle fibers involved in moving chromosomes toward the egg, resulting in “centromere drive”—an advantage in getting into the egg.**
6. **A mutation in a W chromosome that makes it more able to move toward the egg would tend to spread through the population at the expense of W chromosomes lacking that advantage (Fig. 1B).**
7. **Because they are genealogically linked, a sweep in a driving W means a sweep in the mtDNA as well (Fig. 1C).**

**Figure 1.**
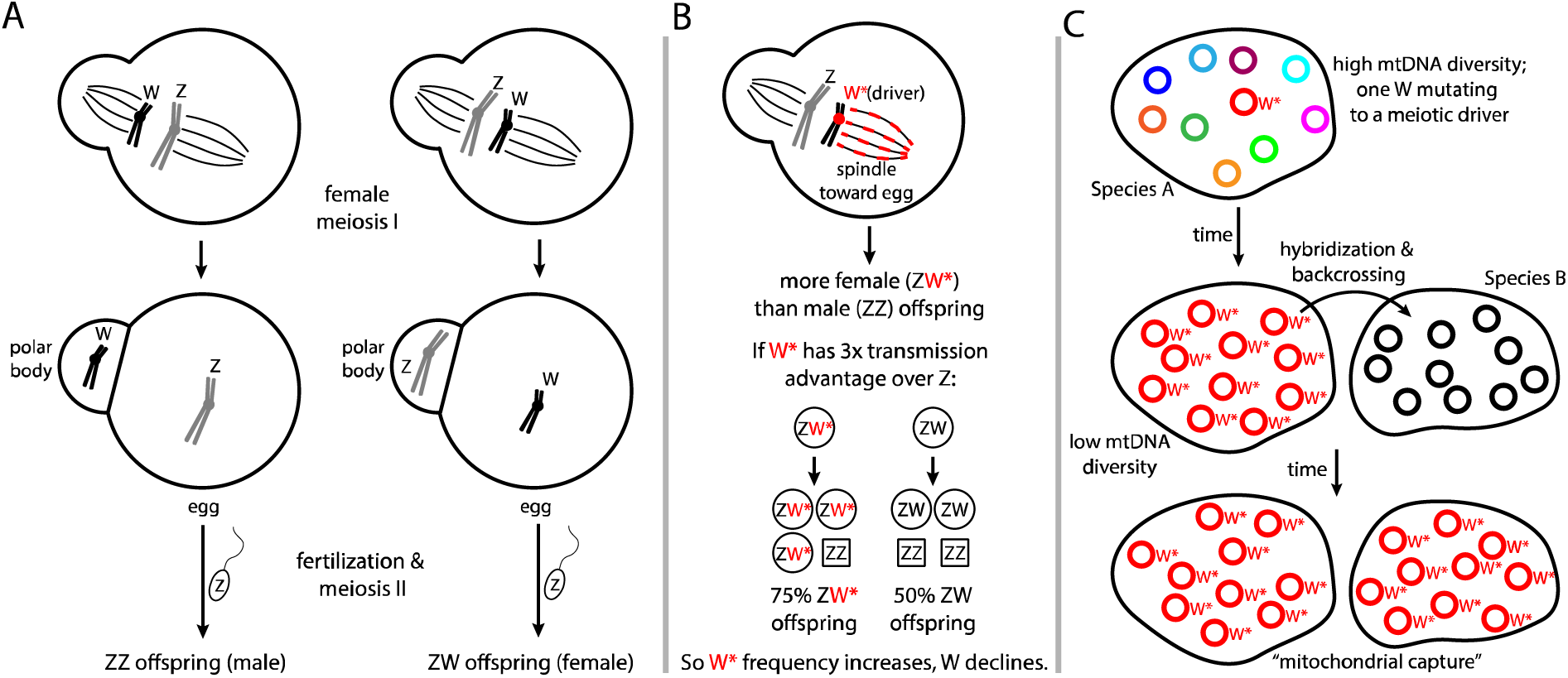
Transmission of the W and Z chromosomes via female meiosis and the potential of a driving W to gain a bias in transmission, resulting in spread of the driving W and associated mitochondrial DNA (mtDNA). (A) Two possible alternatives for the Z and W chromosomes during asymmetric cell division during meiosis I, with one set of spindle fibers leading to the large egg and the other set leading to a small polar body. (B) If a mutant W interacts more strongly with the spindle leading to the egg, it has biased transmission and rises in frequency compared to the non-driving W in the next generation. (C) Jumping up to the level of populations of individuals on a landscape, spread of the driving W carries along the associated mitochondrial type, causing low mtDNA diversity and between-species mitochondrial transfer and replacement (often termed “mitochondrial capture”).

I discuss each of these propositions and the supporting evidence for them below.

Matrilineal mtDNA (#1) occurs in the great majority of eukaryotic species groups (Birky, 2001), although there are some interesting exceptions (Ballard & Whitlock, 2004). ZW sex chromosome systems (#2) occur in many groups, including birds, butterflies and moths, most snakes, and many sub-clades within fish, amphibians, lizards, crustaceans, and plants (Bachtrog et al. 2014; Pennell et al. 2018). In most of these groups, W chromosomes do not recombine with the Z chromosome, except for in the usually small pseudoautosomal regions (Otto et al., 2011).

The genealogical linkage between mtDNA and W chromosomes (#3) has been discussed in the literature (Berlin et al., 2007; Berlin & Ellegren, 2001; Cosmides & Tooby, 1981; Dowling et al., 2008; Gu et al., 2023; Irwin, 2018; Lane, 2008; Marais, 2007), but it has received little attention in the field of mitochondrial phylogeography. Perhaps this is because the W chromosome tends to be more difficult to assemble compared to the Z and autosomes, and as the single-copy sex chromosome is less gene-rich and is often viewed as having less potential for adaptive evolution compared to the Z and autosomes. Despite the W chromosome and mtDNA having different cellular locations and contexts (the single-copy W is in the nucleus and has a role in sex determination and perhaps little else; the mtDNA is in the many-copy cytoplasmic organelles and has a key role in metabolism), their linked inheritance means whatever processes affect the genealogy of one will affect the genealogy of the other in the identical way (with the exception of the pseudoautosomal regions).

The fact that female meiosis of eukaryotes produces a single egg cell (#4) means there is potential competition between differing genetic elements to get into that egg cell. Both cell divisions during female meiosis are asymmetric, with the smaller cell becoming a polar body rather than the egg (Fig. 1A; Dudka & Lampson, 2022; F. Finseth, 2023; Gorelick et al., 2017). There is much evidence for differences in the spindle fibers and other molecular machinery involved in pulling chromosomes towards egg versus polar body (Akera et al., 2017; Kumon et al., 2021).

If chromosomes differ in their ability to interact with the spindle fibers leading to egg versus polar body, they then can differ in their ability to be passed on to the next generation through the egg (#5). The term “transmission ratio distortion” refers to such a situation when one chromosome or allele has an advantage over its counterpart during meiosis. This is also referred to as a form of non-Mendelian inheritance, because it violates our usual expectation that alleles in the nuclear genome have a 50% probability of being transferred to offspring. Despite this, transmission ratio distortion has been observed in female meiosis in a variety of species distributed throughout the tree of life (Clark & Akera, 2021; Didion et al., 2015; Dudka & Lampson, 2022; F. Finseth, 2023; Fishman & Willis, 2005; Henikoff et al., 2001). This can be viewed as a manifestation of adaptation due to selection based on competitive interactions, except that the evolving entity is the chromosome and the competition is between homologous chromosomes for getting into the eggs of the next generation. The competition is intense, as chromosomes in polar bodies leave no descendants (except in very rare cases; Gorelick et al., 2017). The centromere is the chromosomal region that appears most attached to the spindle fibers during cell division—hence the name “centromere drive” for the phenomenon by which some chromosomal variants have an advantage in moving into the egg. The advantage of driving chromosomes over non-driving chromosomes during female meiosis can range from modest to massive; for example, a driving autosome in the monkeyflower *Mimulus guttatus* has a 58:42 transmission advantage against the conspecific non-driving homologous chromosome, and a 98:2 advantage against a heterospecific homologous chromosome in F1 hybrids with *Mimulus nasutus* (Fishman & Saunders, 2008; Fishman & Willis, 2005).

Much progress is being made in understanding the complex molecular interactions involved in centromere drive—for example see Akera et al. (2017); Clark & Akera (2021); F. Finseth (2023); Kruger & Mueller (2021); and Kumon et al. (2021). Briefly, centromeric DNA tends to consist of many repeats of a particular sequence, and known mechanisms of drive tend to involve differences in both DNA sequence and number of repeats. The repeats are wound around protein called centromeric histones, and together the condensed DNA and histones are called heterochromatin. During cell division the heterochromatin interacts with additional proteins to from the kinetochore, the structure that links to the spindle fibers to the centromere. Thus the connection of a chromosome to the spindle fibers can be viewed as an interaction between the DNA sequence (and its repeats), the centromeric histone and kinetochore proteins (which may be coded by genes on other chromosomes), and the characteristics of the spindle fibers (which have a directional signal toward the egg). A mutation in the DNA sequence and/or repeat number can create an advantage for a chromosome in getting into the egg, whereas changes in histone or kinetochore proteins can suppress that advantage (F. R. Finseth et al., 2021; Kumon et al., 2021). There is also some evidence of mutations outside of the main centromeric region that can affect the probability of transmission of chromosomes into the egg (Axelsson et al., 2010; Didion et al., 2015).

Chromosomal mutations that cause meiotic drive advantages have particularly interesting consequences if they occur on W or Z chromosomes. In a single female meiosis, only one of these chromosomes normally succeeds in getting into the egg. A female with a driving W tends to produce an increased proportion of W eggs compared to Z eggs, resulting in a female-biased sex ratio (driving Z would have the opposite result). Although W chromosomes do not find themselves competing against another W in the same cell, they do compete on a population level. A driving W will tend to cause the female they are in to produce more female offspring compared to an individual without a driving W, such that the population frequency of the driving W will increase in relation to the non-driving W (#6; Fig. 1B).

#7 is a logical consequence of #3 and #6. In ZW systems with matrilineal mitochondrial inheritance, surprising genealogical patterns in the mtDNA (e.g., low diversity within and between species) could result from the fixation of a driving W (Fig. 1C).

The plausibility of the Driving W Hypothesis is supported by the evidence in support of each of the seven components above. However, two potential reasons to doubt the potential for fixation of a driving W chromosome are that (1) the Z chromosome might itself evolve drive, counteracting the W drive, and (2) autosomal suppressors of drive might arise. Malik (2009) noted that in ZW sex determination systems, “competition between the sex chromosomes for inclusion into the egg would lead to skewed sex ratios and threaten the population” (see also Hamilton, 1967) and predicted that “any suppressor alleles in autosomal proteins that could alleviate the deleterious effects of this meiotic drive would be selectively swept through this imperiled population.”

To evaluate the relative potency of W drivers compared to Z or autosomal drivers, and to evaluate the potential for autosomal suppressors in stopping W drive, I present simulations.

## Simulation methodology

We can use computer simulations to evaluate the potential for driving W mutations to lead to W-mtDNA sweeps through populations, as well as the potential for suppression of such sweeps. Much of the literature on centromere drive predicts the rise of suppressors that reduce or eliminate the drive effect (Dudka & Lampson, 2022; Henikoff et al., 2001; Malik, 2009; Searle & Pardo-Manuel de Villena, 2024), but there has been little quantitative modelling of the potential for this, especially with regard to W drivers.

To conduct simulations, I wrote scripts in the Julia programming language (Bezanson et al., 2017). Full scripts and explanation are available at https://darreni.github.io/DrivingW, and a summary of the logic of the simulations is as follows.

A population of diploid individuals has constant population size of *N* = 20,000 and non-overlapping generations. There are two major types of sex chromosomes, referred to as Z and W. Each individual carries two sex chromosomes, with ZW individuals being female and ZZ individuals being male. The initial sex ratio of the population is 1:1. Each individual also carries two autosomes. In subsequent generations, each offspring’s parents are determined through a random sample of one female and one male in the parental population.

The three categories of chromosomes (Z, W, and autosome) each can vary in two traits. These are the meiotic attraction strength (*m*) to the spindle fibers leading to the egg, and the ability to suppress meiotic drive (*s*). Hence a focal chromosome *i* has spindle attraction *m_i_* and suppression strength *s_i_*. In the simple case of a chromosome with no meiotic drive ability and no suppression strength, *m_i_* = 1 and *s_i_* = 0. In the simulations presented below, cases are included in which some Z, W, and/or autosomes have a spindle attraction advantage (*m* > 1), and in which some autosomes have an ability to suppress such drive (*s* > 0).

The inheritance of chromosomes from a female parent to offspring is determined by a random draw with probabilities adjusted to reflect the spindle attraction and suppression strength of that parent’s chromosomes. First, the drive proportion (*d_i_*) of a focal chromosome *i* is determined as

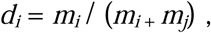

where *m_j_* is the spindle attraction of the other chromosome in the pair. When two chromosomes have equal spindle attraction, the above formula results in *d_i_* = 0.5. Next, overall drive suppression strength (*S*) is calculated as the average of the suppression strengths of the two autosomes, designated as *s_u_* and *s_v_*:

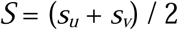

Then, the two factors above are combined to determine the probability (*t*) of a focal chromosome (*i*) being transmitted from mother to offspring:

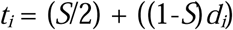

The above way of calculating the transmission probability (*t_i_*) means that when *S* = 0, meaning no suppression, then the transmission probably equals the drive proportion (*t_i_* = *d_i_*). At the opposite extreme, when *S* = 1, there is complete drive suppression (*t_i_* = ½).

For each of the male parent’s chromosomes, transmission probability to the offspring is 0.5 (that is, unbiased random segregation).

Simulations were run with drive strengths (*m*) of 1.1 (a 10% advantage of getting into the egg), 1.5 (50% advantage), or 3.0 (200% advantage) compared to non-driving chromosomes (which have drive strength of 1.0). At the start of most simulations, the frequency of the driving chromosome is 5%, with the rest of the chromosomes being non-driving.

Drive suppressors were incorporated into two simulations with the purpose of inferring the potential for suppressors to slow or prevent the spread of W drivers. In the first, 5% of the starting females have a W driver with *m* = 3.0 and another 5% of the females and 5% of the males have two copies of an autosome with a partial drive suppressor (*s* = 0.5). The second was like the first except the drive suppressor was raised in strength (*s* = 1.0) such that it completely suppresses drive when the suppressor is homozygous.

Note that the values used here for drive strength are well within the range of known empirical examples of female meiotic drive, for example the 98:2 advantage observed for an autosome within hybrids of the monkeyflowers *Mimulus guttatus* and *Mimulus nasutus* (Fishman & Saunders, 2008; Fishman & Willis, 2005) and an up to 95:5 advantage observed for an autosomal locus in house mice *Mus musculus* (Didion et al., 2015). Strengths of suppressors are less well known, but they are regularly invoked in the theoretical literature as having the potential to completely counter the effects of drive (Henikoff et al., 2001; Malik, 2009). Hence the simulations encompass biological plausible scenarios given our current understanding.

I also built a deterministic version of the model, in which population size is assumed to be large enough that it can be treated as infinite. This allows computational efficiency because only relative proportions of individual genotypes need to be tracked, rather than having to track all individuals. It also means that there is no variation between runs, meaning genetic drift has no role. All details of mating, transmission advantage, and suppression remain the same as in the individual-based model. Using this deterministic model, I modelled the cases of 5% driving W chromosomes, 5% driving Z chromosomes, simultaneous 5% driving W and 5% driving Z, and 5% driving W and 100% driving Z (i.e., initially fixed), with all driving chromosomes having *m* = 1.5. All starting populations had combinations of chromosomes according to Hardy-Weinberg equilibrium and linkage equilibrium (i.e., assuming independent combinations of chromosome types, given their population frequencies).

## Simulation results

These simulations provide insight regarding several key questions:

### What is the relative potential of selective sweeps for equivalent-strength W, Z, and autosomal drivers?

Simulations reveal that W drivers (Fig. 2, top row of panels) tend to spread much faster through the population than Z drivers (Fig. 2, middle row) that have equivalent advantage in getting into the egg during female meiosis. Autosomal centromere drivers (Fig. 2, bottom row) have an intermediate speed of spread. These different speeds of spread for equivalent strength drivers can be understood to be a result of the proportion of each type of chromosome that were exposed to female meiosis (versus male meiosis) in the parents of each generation: all W chromosomes, 1/3 of Z chromosomes, and ½ of autosomes (assuming equal sex ratios in the parents). Proportionally then, W chromosomes have experienced meiotic competition to get into the egg 3 times more often than the Z, and 2 times more often than autosomes. Selection for meiotic drive is more intense for the W, making fast spread of W drivers quite likely. (See Hamilton [1967] for similar reasoning applied to Y drivers in the case of male meiosis.)

**Figure 2.**
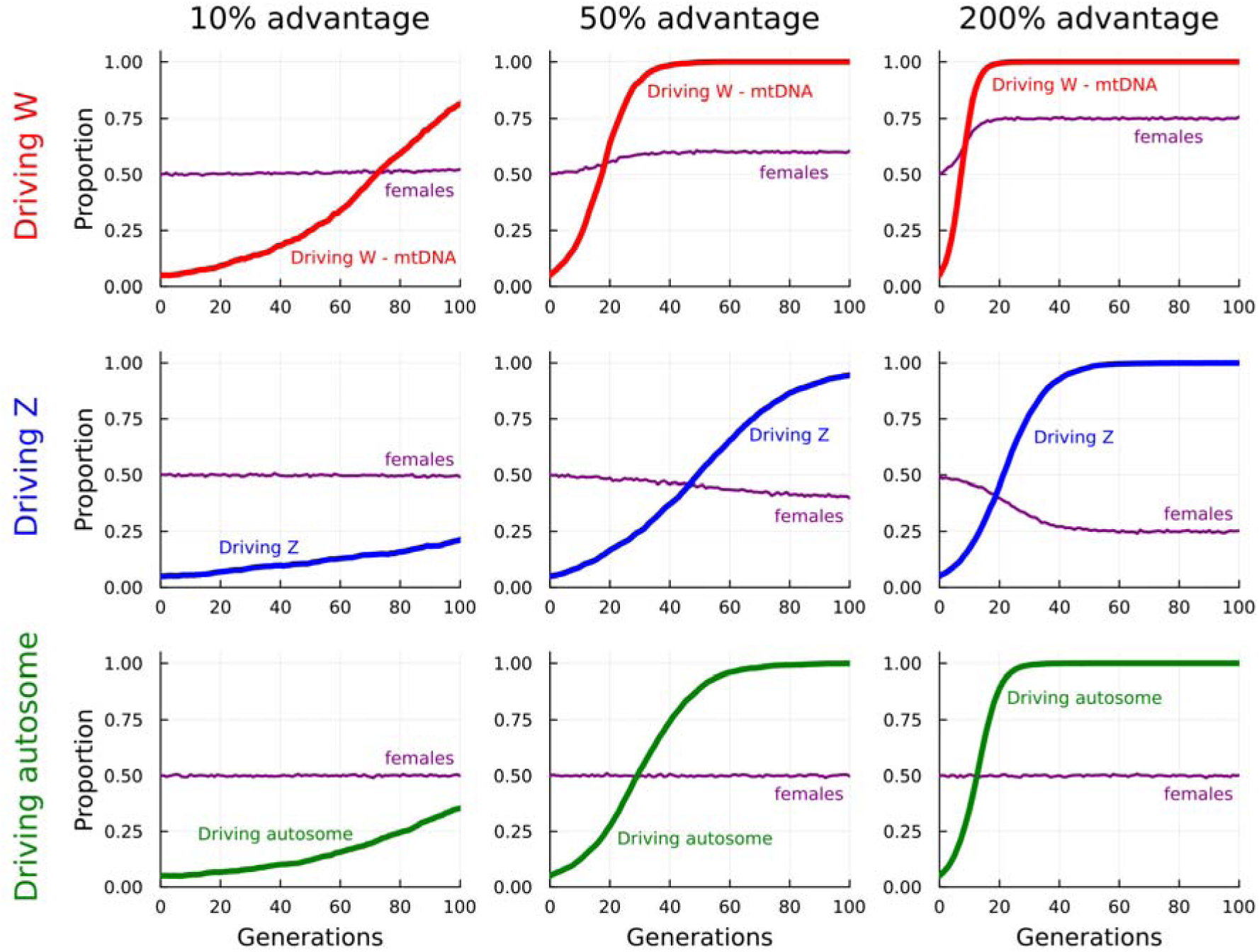
Driving W chromosomes (and their associated mtDNA) spread more quickly through population than equivalent strength driving Z chromosomes or autosomes. Each panel shows growth in population frequency of a driving chromosome: either a driving W (top row of panels); a driving Z (middle row); or a driving autosome (bottom row). These are from separate simulations in which each driver starts at a frequency of 5% and has either 10% (*m* = 1.1; left column), 50% (*m* = 1.5; middle column), or 200% (*m* = 3.0; right column) greater attraction to the egg compared to a non-driving chromosome (for which *m* = 1.0). Population size is 20,000 and mating is random. The proportion of females (ZW individuals) is shown by purple lines.

### Do evolutionarily important W drivers necessarily have a large impact on the sex ratio?

A driving W with a 10% advantage in getting into the egg rises from 5% frequency to fixation in about 200 generations (short in evolutionary time) and results in a sex ratio skew of only about 0.5238 female to 0.4762 male (i.e., 10% more females compared to males), a difference that would be difficult to detect in many natural study systems (see upper left panel of Fig. 2). This is a manifestation of the general challenge of studying selective forces in natural populations: a small selective advantage over many generations can have massive impacts on allele frequencies of a population yet be difficult to detect in any one generation. Another reason that spread and fixation of W drivers might not result in much sex ratio skew is that subsequent spread of Z drivers or autosomal suppressors can move the sex ratio back towards 1:1.

### Does a Z driver reduce the potential of a W driver to spread?

No. When W and Z drivers of equivalent strength start out simultaneously at low frequency, the spread of the driving W is not slowed by the (much slower) spread of the driving Z (Fig. 3A-C). When a Z driver is already fixed, then the W driver tends to spread slightly more quickly through the population (Fig. 3D). This is because when the sex ratio is not 1:1, alleles that cause a higher proportion of the rare sex tend to have an advantage (Shaw & Mohler, 1953). This reinforces a point made above, that while the W and Z can be viewed as competing to get into the egg, at a population scale the W is competing with other W’s, and the Z with other Z’s. Hence W drivers and Z drivers tend to encourage the spread of each other, causing a tendency for the sex ratio staying close to 1:1 (Fig. 3C,D).

**Figure 3.**
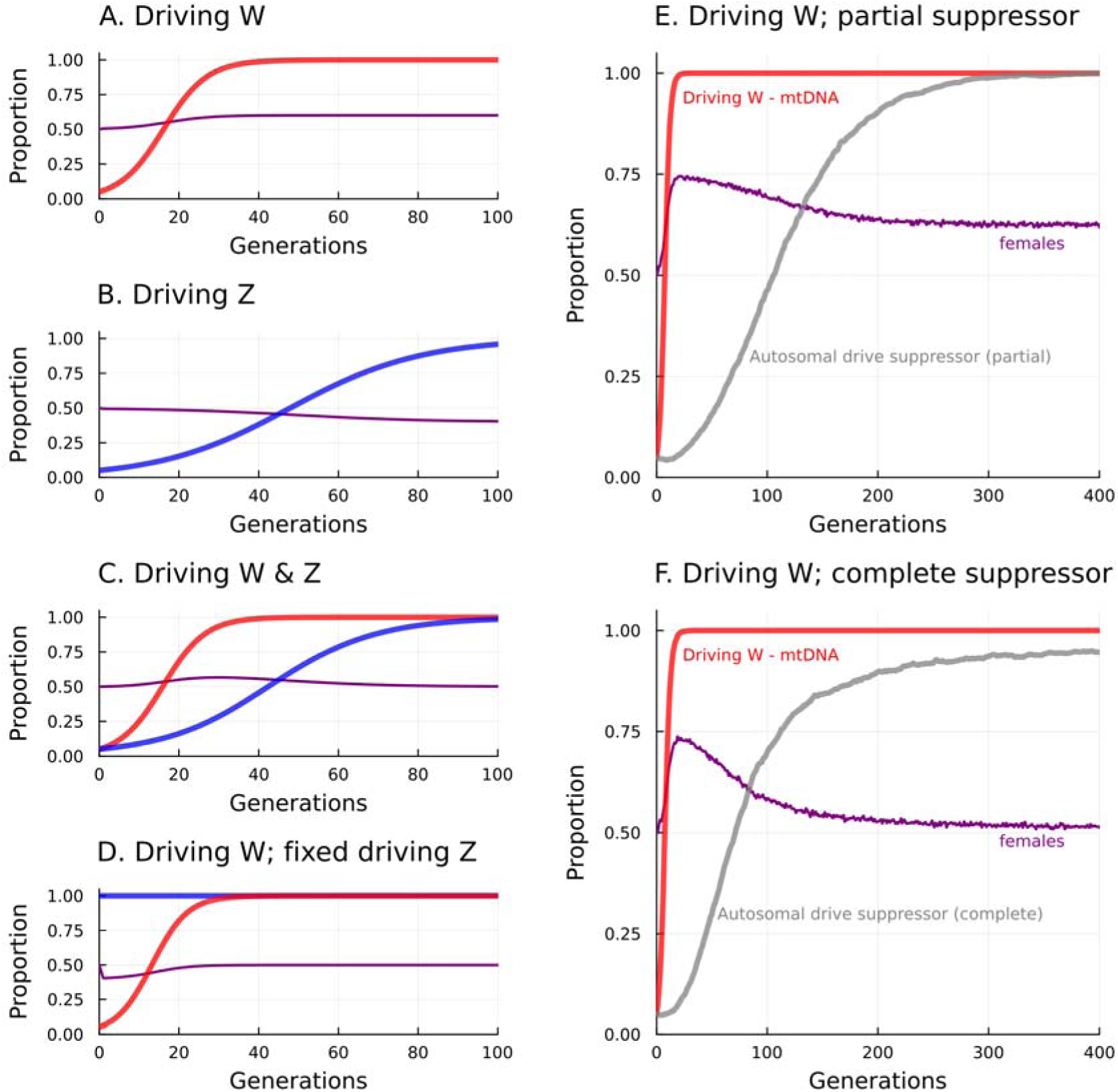
A driving W chromosome spreads quickly to fixation (along with the associated mtDNA), regardless of driving Z or autosomal suppressors that arise around the same time. Each panel shows the growth in frequency of a driver (W in red, Z in blue) and/or suppressor (in grey) from 5% initial frequency, with female proportion shown in purple. In panels A-D, drivers have 50% greater attraction to the egg (*m* = 1.5) compared to non-drivers (*m* = 1.0); in panels E-F, drivers have 200% greater attraction to the egg (*m* = 3.0) than non-drivers. Panels A and B show the spread of W and Z drivers on their own, whereas C shows their simultaneous spread, and D shows the W driver against an already fixed Z driver. Panels E and F show that the growth of autosomal repressors of W drive is slow compared to the spread of the W driver. In panel E, the suppressor (*s* = 0.5) reduces the W drive advantage by half (when homozygous; one quarter when heterozygous) in an individual with the W driver, whereas in panel F, the suppressor (*s* = 1.0) completely eliminates the W drive advantage (when homozygous; half suppression when heterozygous). In A-D, simulations are done deterministically (so no stochasticity due to limited population size), whereas in E-F population size is 20,000. Mating is random.

### How much potential is there for novel autosomal suppressors to prevent spread of W drivers?

In simulations in which a driving W and autosomal suppressor start out at the same low frequency, the driving W spreads much faster than the suppressor (Fig. 3E,F). This is for two reasons. First, when the W driver is at low frequency, there is strong selection for the W driver but there is virtually no selection for the suppressor because the population sex ratio is still close to 1:1 (A. W. F. Edwards, 2000; Shaw & Mohler, 1953; Shyu & Caswell, 2016). Second, because the driving W and autosomal suppressor segregate independently, in the early phase of the spread of a W driver the suppressor is only rarely in the same pre-meiotic cell as the driver. Only when the W driver spreads enough to produce a strongly skewed population sex ratio does the selection for the suppressor become strong. Even then though, the suppressor finds itself in male meiosis (where it experiences no advantage) about half of the time, whereas the W driver is always in competition to get into the egg. The driving W goes to fixation on a vastly shorter timescale than the suppressor (Fig. 3E-F). This is true for both complete and partial suppressors; while a complete suppressor has a higher maximal rate of spread than a partial suppressor (compare steepest slope of grey line in Fig. 3F with that in Fig. 3E), a complete suppressor is ironically less likely to go to full fixation in a fixed period of time than a partial suppressor. This is because as a complete suppressor causes the population sex ratio to get closer and closer to 1:1, the selective advantage of the complete suppressor approaches zero (Shaw & Mohler, 1953). In contrast, a biased sex ratio is maintained when there is only a partial suppressor, preserving selection for the partial suppressor even when it is at high frequency (compare female frequency [purple] lines in Fig. 3F and Fig. 3E). The net effect is that both complete and partial autosomal suppressors of W drive are quite slow in their spread, but at differing parts of their expansion curves, and both are vastly slower in their spread compared to a driving W.

## Relationship between drive strength and selective strength

In the above simulations, meiotic drive strength is expressed in terms of *m*, the attraction of a chromosome to spindle leading to the egg, where “normal” non-driving chromosomes have *m* = 1 and a driving chromosome has *m* > 1. I derived formulae for the relationship between *m* and the strength of selection as it is often expressed in the literature, as a selection coefficient. Here I express the selection coefficient as the symbol *c*, which means the additional fitness of a driving chromosome relative to a fitness of 1 for a non-driving chromosome. For a W driver, this is

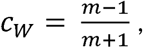

which can be understood as the ratio of the increased spindle attraction of the W driver compared to the total spindle attraction of the W driver and Z chromosomes. So for example, a 50% W drive advantage (*m* = 1.5) results in a selective advantage of *c* = 0.5 / 2.5 = 20% over “normal” W chromosomes (so relative fitness of the driving W is 1.2, compared to 1 for the normal W).

For Z drivers, if we assume Hardy-Weinberg equilibrium and the same frequency of the driver in females and males, the selection coefficient is

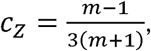

so one-third of the W case, which can be understood as a result of the Z chromosomes spending only 1/3 of the time in females, in a genealogical sense. So in the example of a 50% drive advantage, a Z driver has a selective advantage of only *c* = 0.5 / (3 × 2.5) = 6.67%. Note that these calculations are in terms of the increase in frequency of the driving Z between the parental population and the whole offspring population. In multi-generation simulations, the driving Z in mothers causes an increase in frequency of the driving Z in sons (not daughters, as they receive a W from their mother), and this higher frequency in sons is passed on to the granddaughters of the original mothers. Hence in a realistic case in which the frequency of the Z driver is a little different in males and females, the translation of meiotic drive strength into fitness is slightly more complex than the formula above.

The formula for autosomal drivers is a bit more complicated, because the strength of selection depends on the relative frequency of the driving autosome (*p*) compared to the homologous non-driving autosome. Nonetheless, the form of the relationship between spindle attraction and the selection coefficient is reminiscent of the above:

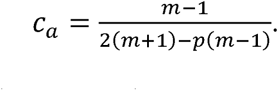

When the driving autosome is rare (*p* near zero), the formula shows that the selection strength is one-half that of a driving W, because the autosome spends half of its time in females. When the driving autosome becomes common, then the strength of selection increases because the non-driving autosome spends a higher fraction of its time in a heterozygous state, exposing it to more of a drive disadvantage. For our example of a 50% drive advantage, the selective advantage ranges from 10% when the autosomal driver is rare to 11.1% when the driver is common.

These considerations show that for mutations that give chromosomes a particular increase in ability to get into the egg, the resulting selection advantage of W drivers is 3 times that of Z drivers, and 2 times that of autosomal drivers that are rare.

## Discussion

In female meiosis in ZW systems, the Z and W chromosomes face stark competition: only one gets into the egg (Fig. 1A,B). In many species, much of the length of these chromosomes ceased recombining with each other a long time ago (e.g., in birds, on the order of 100 million years ago), meaning they have highly divergent sequences and are expected to differ much in their biochemical interactions. Given this, it is remarkable that their ability to move to the egg is roughly equal in most times in most species, leading to the close to 1:1 sex ratios that are usually observed. Rutkowska & Badyaev (2008) have pointed out that given the large differences in the Z and W, there must be evolved mechanisms that prevent transmission bias. Given this, we can view the close to 1:1 ratios typically observed as being the result of a long history of both Z and W evolving opposing mechanisms that provide advantage in getting into the egg and autosomal loci evolving suppressors of those advantages. In fact, there is evidence that many female birds can adjust sex ratio in response to environmental conditions (Komdeur, 1996; Rutkowska & Badyaev, 2008; Sheldon, 1999), suggesting physiological features that can be manipulated to affect whether the Z or W moves into the egg. Given this evidence for physiological influences on Z or W movement into the egg, it is reasonable to think changes in the Z or W can influence that as well. The logic and simulations presented above indicate more sweep potential for W drivers than Z and autosomal drivers of equivalent drive strength, and little potential for suppressors to prevent a W sweep.

Is there empirical evidence that such W-driven sweeps of mtDNA are happening regularly in nature? In fact, there are intriguing observations consistent with the W-mtDNA drive hypothesis. Within-species diversity at the W chromosome is often dramatically lower than at autosomes, for example 8-13 times lower in flycatchers (Smeds et al., 2015) and 13-28 times lower in chickens (Berlin & Ellegren, 2004). These differences are far larger than the expected 4 times difference based on the different inheritance modes of W chromosomes and autosomes. Despite the low within-species diversity of the W, it undergoes rapid between-species structural evolution in birds (Rutkowska et al., 2012; Sigeman et al., 2024), snakes (Augstenová et al., 2018), and butterflies and moths (Han et al., 2024). These observations show that the low within-species diversity of the W is not due to lack of mutational change in the W; rather, the picture emerges of a rapidly evolving chromosome that nonetheless has little within-population variation. Moreover, changes in repetitive element number and location are involved in these cases of rapid W evolution, which is intriguing because repetitive elements tend to be associated with centromeres and known cases of centromere drive (F. Finseth, 2023; Malik, 2009). These observations are consistent with structural mutations in the centromeres causing repeated rounds of centromere drive, leading to regular sweeps of the W and the associated mtDNA through a population, and occasionally between populations. I note also that in plants with ZW sex chromosome systems, W drive is expected to influence chloroplast DNA (cpDNA) variation as well as mtDNA, since cpDNA is usually maternally inherited.

A key observation consistent with the driving W hypothesis is that within-population mtDNA variation (synonymous site diversity) is on average roughly threefold lower in birds, which have ZW sex chrom osomes, than in mammals, which have XY sex chromosomes (Berlin et al., 2007). Moreover, Berlin et al. reported two other statistics (the ratio of nonsynonymous to synonymous diversity; and the neutrality index) that differed strongly between birds and mammals, in the direction consistent with birds having reduced efficacy of selection against deleterious mutations on the mtDNA. Berlin et al. (2007) interpreted these differences as being due to Hill-Robertson effects on the W chromosome, in which selection at closely linked sites causes a reduction of effective population size and thereby results in low diversity and less effective purging of deleterious mutations. While not mentioned by Berlin et al., mutations that cause meiotic drive can be viewed as a related phenomenon: a driving W chromosome (in birds) would cause sweeps in both W and mtDNA, lowering diversity and reducing the impact of selection against deleterious mitochondrial variants. There are however other possible reasons the effective population size might be lower in birds than mammals. To investigate further, incorporating measures of autosomal diversity into this analysis would help control for potential differences in population size between birds and mammals. Ultimately though, we need more comparisons between independent ZW and XY clades, of which there are many throughout the tree of life (Pennell et al., 2018; Wilson Sayres, 2018), to more effectively test whether differences in mitochondrial diversity are due to the link with W chromosomes.

A causal connection between the W chromosome and low mtDNA diversity is suggested by the above evidence of (a) extremely low within-species W-mtDNA diversity compared to autosomal diversity and (b) lower mtDNA diversity in ZW compared to XY systems. How can we distinguish between meiotic drive or regular whole-organism natural selection (not involving meiotic drive) on the W as being the cause? First, W drive will cause immediate within-family offspring sex-ratio distortion that could in principle be detected at the single-cell zygote stage. In contrast, survival-based natural selection on the W might cause some sex ratio distortion but this would take time to develop over the life spans of the offspring. Second, W drive will tend to eventually lead to the spread of autosomal or Z-chromosome suppressors of drive—these may be detectable via population genomic analysis of levels of variation across the genome as well as close examinations of variation in genes related to the centromeric histones and kinetochore proteins. Third, W drive might cause some hitchhiking of deleterious W and mtDNA mutations, lowering whole-organism fitness compared to a scenario only involving natural selection. Such decrease in whole-organism fitness might be detected when there is introgression of W-mtDNA between populations due to a W driver. Finally, relative support for the meiotic drive vs. non-drive natural selection scenarios could be evaluated by considering the degree to which each is expected to differ between W chromosomes and other types of chromosomes (Z, Y, X, and autosomes) and then comparing empirical data to those expectations. As shown by the simulations, female-based meiotic drive is expected to be especially powerful on the W compared to the Z and autosomes (and does not apply to the Y because it does not experience female-based meiotic drive). In contrast, natural selection applies to all of these chromosomes but to varying degrees based on factors such as gene content and recombination landscapes. Hence comparisons of diversity of W chromosomes (in ZW systems) and Y chromosomes (in XY systems), where natural selection might be expected to be of similar magnitude but female meiotic driving would apply only to W chromosomes, may be revealing in evaluating support for or against the driving W hypothesis.

Catching a driving W mid-sweep may be challenging, given that W drivers may spread through the population quickly (Fig. 3) and may not affect sex ratio very much, especially at the start of the drive or after a Z driver or autosomal suppressor has returned the ratio to close to 1:1. Perhaps the best place to notice W drivers will be in newly formed hybrid zones, where the spread of a W driver from one population to another might be happening, thereby skewing sex ratios there. Whether a W driver spreads, bringing the associated mtDNA along with it, is of course determined not just by the driver strength but also by potential costs of deleterious mutations on the W or the mtDNA. It is possible that in hybrid zones, the two populations might have differentially adapted mtDNA (perhaps due to mitonuclear coevolution; Hill, 2019; Wang et al., 2021), such that a driver would spread through one population but not into the other. It is also possible that hybrids might have low fitness due to incompatibilities generated as a result of the evolution of different centromere drivers and suppressors in the two populations (Henikoff et al., 2001; Searle & Pardo-Manuel de Villena, 2024; Talbert & Henikoff, 2022). In the case of W drivers and autosomal suppressors that affect only the W drivers, this phenomenon would only manifest in females, contributing to the well-known phenomenon known as “Haldane’s rule” in which the heterogametic sex tends to have lower fitness than the homogametic sex (Frank, 1991; L. D. Hurst & Pomiankowski, 1991).

In the simulations presented here the only potential whole-organism cost to the W drivers is the biased population sex ratio, and we have seen above that autosomal suppressors spread slowly compared to the driver. In the literature a variety of other potential costs are discussed (Kumon & Lampson, 2022; Malik, 2009). One is that the driver could spread deleterious variants that are physically linked to the driving allele. In the case of W drivers, that applies also to genealogically linked deleterious variants on the mtDNA as well as the W. The potential power of a W driver to spread would then be jointly determined by its drive advantage and the costs of these deleterious effects at genealogically linked loci. Such deleterious effects may accumulate especially rapidly on large non-recombining genomic regions such as the W chromosome; however this is true for all copies of the W in the population, and the mutation of a non-driving W to a driving W might occur on a W with higher or lower fitness costs of such deleterious mutations than the population average. Another potential cost is that centromere drivers could cause problems in male meiosis (Fishman & Saunders, 2008), but that would not apply for a W driver because males do not have a W chromosome. There is also the idea that centromere drivers could cause problems in mitosis, if the driving centromere attracts so many spindle fibers that other chromosomes are not as able to bind them, resulting in imbalanced mitosis (Drpic et al., 2018; Kumon & Lampson, 2022). For W drivers this would again not affect males since they have no W, but mitosis in females could be impacted, reducing fitness in a way that is again consistent with Haldane’s rule (see above). Finally, the literature on meiotic drive often refers to the possibility of a stable polymorphism, when a driver spreads when rare but then stops spreading because of a high fitness cost when in the homozygous state, due to exposed recessive deleterious mutations (Hartl, 1970). This phenomenon also does not apply to the W chromosome because it occurs only in a hemizygous state. Overall, these considerations point to lower potential cost to W drivers than to equivalent-strength Z or autosomal drivers, again pointing to their potential importance in causing sweeps of mitochondrial variants.

While here I have focused on the scenario of a W chromosome experiencing a mutation that gives it a meiotic drive advantage, another possibility was raised by Cosmides & Tooby (1981): perhaps mtDNA could mutate in such a way that creates a sex ratio bias towards females, via an increase in survival of female zygotes compared to male zygotes. This would tend to cause population spread of that mitochondrial variant along with its associated W chromosome. Cosmides & Tooby (1981) commented that the effects of this would be “difficult to distinguish from meiotic drive,” by which they apparently meant drive by a W chromosome. This brings up another intriguing scenario: Might it be possible that a mitochondrial variant could cause a transmission bias of the W into the egg? It is known that mitochondria are highly involved in interacting with spindle fibers during cell division (Scholey 2025), raising the possibility that mitochondrial gene products contribute to preferential interaction between a W chromosome and the spindle fibers leading to the egg. As Cosmides & Tooby (1981) pointed out, the selective interests of the W and mtDNA are completely aligned, hence these kinds of synergistic interactions are plausible.

For technical and theoretical reasons, there has historically been more research focus on the major sex chromosomes (X and Z) compared to the minor sex chromosomes (Y and W), but the latter are nonetheless important in evolution (Mank, 2012). Although there tend be fewer functional genes on Y and W chromosomes compared to many autosomes, those genes can play a key role in sexual selection and speciation (Mank, 2012; Wright et al., 2014). Here, we have seen that the genealogical link between the W chromosome and mtDNA means that driving W chromosomes might have massive influence on the mitochondrial phylogenies that have played such a central role in the fields of phylogeography, speciation biology, molecular ecology, and taxonomy. The W has tremendous potential for recurrent episodes of centromere drive, thereby reducing diversity in the mitochondria within and sometimes between populations. While natural selection based on whole-organism fitness and mitonuclear coevolution are clearly part of the story (Ballard & Whitlock, 2004; Hill, 2019), I predict that the W-mtDNA Drive Hypothesis will contribute much to our understanding of why mitochondrial DNA often shows so little within-population variation, such clear differences between populations, and otherwise unexplained cases of mitochondrial transfer between species.

## Acknowledgments

I thank Jessica Irwin, Sally Otto, Finola Fogarty, Claudie Pageau, Caleigh Charlebois, Graeme Keais, Diala Abu Awad, Max Reuter, and four anonymous reviewers for helpful discussion and comments on the manuscript. I note that sex chromosomes and sex roles are remarkably complex and often fluid in the biological world (Gorelick et al., 2017; Pennell et al., 2018), and the simplifications that are made for the purposes of building useful models for the study of evolutionary dynamics should not be taken as meaning that sex and gender are binary.

Although not applicable to a theory and simulation paper, I confirm: All the research complies with applicable laws on sampling from natural populations and animal experimentation (including the ARRIVE guidelines).

## Data Availability Statement

All simulation and graph-making scripts (in the Julia programming language; Bezanson et al., 2017) are shown at https://darreni.github.io/DrivingW.

